# Influence of contextual exposure on memory strength and precision for inhibitory avoidance in male and female rats

**DOI:** 10.1101/2025.02.27.640186

**Authors:** Aspen R. Holm, Jason J. Radley, Ryan T. LaLumiere

## Abstract

Aversive associative learning paradigms such as inhibitory avoidance (IA) are frequently used to examine episodic-like memories in rodents. In IA, rodents learn to associate a context with a footshock, followed by testing for memory strength in the original training context and for memory precision in a similar yet distinct neutral context. The present work assessed the effects of different contextual exposure procedures on memory strength and precision in IA at both recent and remote time points using male and female Long-Evans rats. An initial experiment found that rats kept in the lit (non-shock) compartment of the IA apparatus for 60 s during training, as opposed to 10 s, displayed enhanced memory strength, with discrimination between both chambers at the recent retention test and generalization at the remote retention test. Subsequent experiments investigated the effects of contextual pre-exposure the day before training. The results indicate that pre-exposure to the neutral context promoted generalization without altering memory strength compared to the first experiment. In contrast, pre-exposure to the aversive chamber promoted discrimination and enhanced memory strength. Notably, the different procedures yielded similar effects in both sexes. However, the results also indicate an overall pattern of greater contextual discrimination in females compared to males. These findings provide evidence for how different contextual exposures influence the degree of encoding at the time of training and a behavioral foundation for future studies examining the neurobiological mechanisms underlying memory strength and precision in IA, while highlighting the importance of using both sexes in initial behavioral work.

**Significance Statement:** Strength and precision are two fundamental properties of memory that can be simultaneously measured using inhibitory avoidance (IA), a type of context-footshock association task. However, little is known about how different context exposures alter rats’ encoding of these memories, thereby influencing subsequent memory strength and precision. Here, we found that pre-exposure to the neutral IA chamber decreased memory precision, whereas pre-exposure to the aversive IA chamber promoted memory strength and precision. Additionally, females demonstrated overall enhanced memory precision compared to males. These results indicate that different types of contextual exposures influence initial IA encoding and add to a limited body of research examining memory strength and precision in IA in both sexes.

## Introduction

Episodic-like memories exist along at least two dimensions: strength and precision. Memory strength, or the degree of retention for a particular event, is frequently measured using context-footshock associations in rodents (Santos-Anderson and Routtenberg, 1976; Boast and McIntyre, 1977). In these types of learning tasks, animals are given a footshock in a particular context and then tested for their retention in that same context at a later time point. Stimulus generalization can occur as rodents respond to stimuli that are similar to but distinct from the conditioned stimulus (Pavlov, 1927; Guttman and Kalish, 1956). In context-footshock learning, memory precision, or the degree of discrimination versus generalization between similar contexts, has traditionally been assessed by testing animals in both the original training context as well as a contextually modified yet similar context.

Inhibitory avoidance (IA) is one such context-footshock association task that has frequently been used to investigate the neurobiological mechanisms underlying memory consolidation in rodents, with a primary focus on the memory strength for the original learning event (Parent et al., 1995; McIntyre et al., 2002; Huff et al., 2013). However, as recent studies can attest, IA can also be used to assess memory precision (Atucha and Roozendaal, 2015; Met Hoxha et al., 2024).

Previous work suggests that the degree of contextual exposure around the time of training influences retention, presumably due to the extent of contextual encoding. Indeed, prior evidence has found that pre-exposure to the training context plays a critical role in contextual encoding, often leading to enhanced retention, in a variety of behavioral paradigms including IA (Fanselow, 1990; Bae et al., 2015; Huff et al., 2016). However, fewer studies have investigated the effects of pre-exposures on memory precision in IA.

Therefore, to address how different kinds of contextual exposures around the time of training influence IA retention, the current work investigated the effects of several different behavioral procedures during or before training on memory strength and precision. Moreover, as considerable evidence suggests that rodents show increasing context generalization as time passes following the original training, consistent with many theories of systems consolidation (Perkins and Weyant, 1958; Wiltgen and Silva, 2007; Wiltgen et al., 2010; Atucha et al., 2017; Pollack et al., 2018), the present studies also examined retention at both recent (2 d) and remote (28 d) time points after IA training to determine whether these procedures had long-lasting consequences for strength and precision. The first experiment examined how retaining rats in the lit (non-shock) compartment of the IA apparatus for 10 versus 60 s altered subsequent retention measures. A follow-up experiment investigated how testing in the two chambers on the same day, as opposed to alternating days, influenced the results. Two subsequent experiments examined how pre-exposure to the neutral or the aversive training context one day before IA training influenced subsequent measures of strength and retention. As most prior IA studies have used only male rodents, the present work used both males and females to assess whether sex influences the effects of any these procedural differences. Together, the results suggest that neutral context pre-exposure biases rats toward generalization whereas pre-exposure to the aversive context biases rats toward discrimination.

## Methods

### Subjects

Adult male and female Long-Evans rats (10-11 weeks at time of arrival) from Charles River Laboratories (N = 92) were used for these experiments. Rats were single-housed in a temperature-controlled environment under a 12-hour light/dark cycle (lights on at 06:00) and allowed to acclimate to the vivarium for at least 2 days prior to the start of handling. Food and water were available ad libitum during all behavioral experiments. All procedures were compliant with the National Institutes of Health *Guidelines for the Care of Laboratory Animals* and were approved by the University of Iowa Institutional Animal Care and Use Committee.

### Behavioral procedures/apparatus

#### Apparatus

All rats were trained on a single-trial step-through IA task and were tested in both the original context (hereafter, the “aversive context”) and a slightly modified neutral context. The IA chambers each consisted of a trough-shaped box (91 cm long and 20 cm deep) divided into two compartments. The first third was the non-shock compartment, which had white plastic floor and walls and was illuminated by a table lamp, and the second two-thirds comprised the shock compartment, which was made of stainless steel and was not illuminated (Figure 1). The chambers contained a stainless-steel retractable door separating the two compartments. The aversive chamber was connected to a shock generator (Lafayette Instrument Company) controlled by a timer. Both chambers were kept in the same room on tables with approximately 1.25 m separating them. The neutral context was modified to appear contextually distinct in the following ways: 1) The dark compartment contained eight 2-inch wide pieces of white tape on the walls and floor of the dark compartment; 2) The lit, non-shock compartment of the aversive chamber was made of smooth plastic, whereas the non-shock compartment of the neutral chamber was made of textured plastic; and 3) The neutral chamber was cleaned between each rat with a lemon-scented multipurpose cleaner, whereas the aversive context was cleaned with a 20% ethanol solution between each rat.

**Figure 1.**
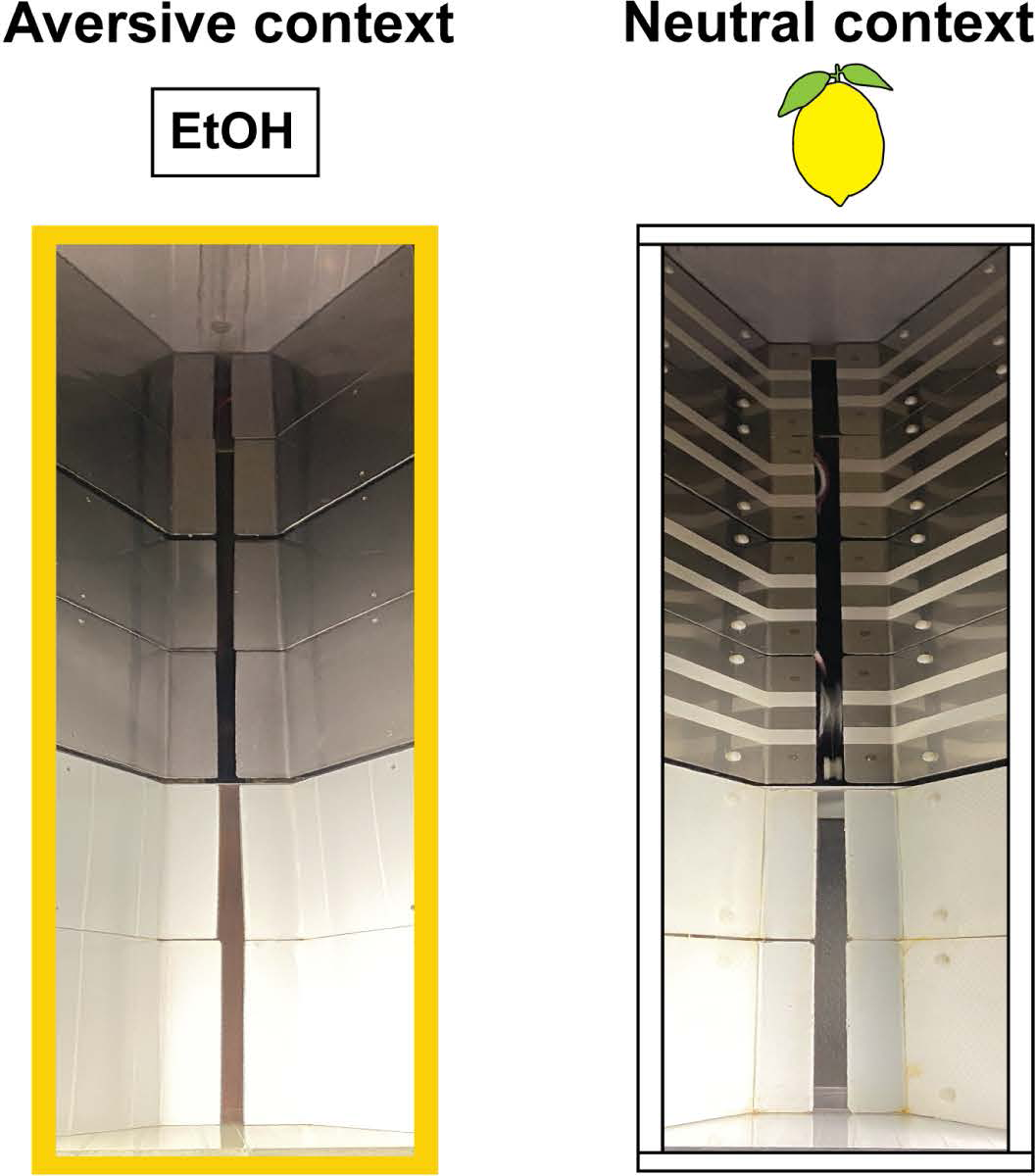
Pictures of IA chambers, with the aversive chamber shown on the left and the neutral chamber shown on the right.

#### General IA procedures

Prior to any behavioral training, all rats underwent 3 d of handling, wherein they were handled individually for 1 min per day. Rats were handled in a room separate from the room in which they underwent behavioral training/testing and were allowed to acclimate to the handling room for 30 min before and after handling each of the 3 days. Additionally, during all handling, training, and testing, white noise (60 dB) was played.

For training and testing, rats were kept in their homecages in the handling room for 1 h before and after training and testing. During IA training, rats were placed in the lit compartment of the aversive IA chamber facing away from the retractable door, which was in place to separate the two compartments. Rats were left in the lit compartment for either 10 s or 60 s (see Experiment 1), the door was lowered, and rats were allowed to cross into the dark compartment. When the rat crossed over with all four paws, the door was raised, forcing the rat to remain in the dark compartment. After 20 s, the rat received a single inescapable footshock (0.5 mA for 1 s). The rat remained in the dark compartment for another 20 s following footshock, after which the rat was removed and returned to its homecage.

Rats were tested for their retention 2-3 d (recent) and 28-29 d (remote) later, with testing occurring in the aversive and neutral context on separate days in a counterbalanced manner, except for Experiment 2 in which testing in both chambers occurred on the same day. During testing, each rat was placed in the lit compartment of each chamber facing away from the retractable door. After 10 s in the lit compartment, the retractable door was lowered, and the rat was allowed to cross into the dark compartment. The latency to cross into the darkened compartment with all four paws was used as the measure of retention in both chambers, with a maximum latency of 600 s.

#### Experimental design - Experiment 1: 10 s versus 60 s holding period

Prior work with IA has typically found a large positive skew in the retention test data, with a large cluster of data points showing relatively low retention latencies, and fewer data points approaching higher latencies (Wahlstrom et al., 2018; Glickman et al., 2024). One possible explanation for this skew is that it reflects a wide degree of variation among rats for encoding the context during training. Therefore, Experiment 1 examined whether providing a greater opportunity to encode the context during training would enhance retention and reduce skewness. In Experiment 1, to test this hypothesis, rats were held in the lit compartment during training for either 10 s or 60 s prior to the door being retracted. Otherwise, all procedures were identical to those described in the General IA Procedures.

#### Experimental design - Experiment 2: Same-day testing

The results of Experiment 1 led to all rats in Experiments 2-4 being trained using the 60 s holding period. Additionally, because prior work found differences in discrimination in the IA task when rats are tested in both chambers on the same day versus on different days (Atucha & Roozendaal, 2015), Experiment 2 examined whether testing rats in both chambers on the same day for both recent and remote retention tests, as opposed to the alternating days in Experiment 1, would alter retention measures. Rats were tested in one chamber and then immediately tested in the other chamber before being returned to their homecage. Testing chamber order was counterbalanced. The results of this experiment led to subsequent experiments using the same alternating day testing procedures as Experiment 1.

#### Experimental design - Experiment 3: Pre-exposure to the neutral context

Experiment 3 examined the effects of giving rats a pre-exposure to the neutral context. The day before training, rats were placed in the lit compartment of the neutral context with the retractable door open and were allowed to freely explore the entire chamber for 5 min. The next day, rats underwent IA training with a 60 s holding period as described above. Rats were tested for retention 2-3 d and 28-29 d after training, with testing in each chamber on a separate day, as described in Experiment 1.

#### Experimental design - Experiment 4: Pre-exposure to the aversive context

Experiment 4 examined the effects of giving rats a pre-exposure to the aversive chamber. The pre-exposure procedure was the same as in Experiment 3, except rats were placed in the aversive chamber instead. The training and testing parameters were the same as in Experiment 3.

### Statistical analysis

GraphPad Prism 10 was used for all analyses. Retention test latencies were analyzed using two-way repeated measures ANOVAs matched by either the two test environments (aversive and neutral), the two retention tests (recent and remote), or matched by both factors, or one-way between-subjects ANOVAs. Post-hoc pairwise comparisons were performed using the Holm-Sidak multiple comparisons test. Paired t-tests were also used to analyze changes in memory strength between recent and remote retention tests, and unpaired t-tests of discrimination ratio values [aversive context latency/(aversive context latency + neutral context latency)] were used to analyze discrimination levels. Skewness was calculated for the 10 s and 60 s retention test data in Experiment 1 using Pearson’s second coefficient of skewness [3*(mean – median) / standard deviation]. All retention test data was also analyzed separately for males and females to determine differences in IA retention between the sexes. Data are expressed as the mean ± SEM, and the α level was set at 0.05. Group sizes were based on previous work from our laboratory, and each group’s n is shown in its respective figure.

## Results

### Experiment 1

Experiment 1 examined whether keeping rats in the lit compartment during training for 10 s versus 60 s before door retraction would alter retention measures. Figure 2A shows the experimental timeline. Figure 2B shows the recent and remote retention test latencies in the 10 s (left) and 60 s (right) holding conditions. For rats in the 10 s condition, a two-way repeated-measures ANOVA revealed no significant main effect of recent versus remote test (F_(1, 18)_ = 1.90, *p =* .19), no significant main effect of context (F_(1, 18)_ = 2.61, *p =* .12), and no significant interaction (F_(1, 18)_ = 0.05, *p =* .82). Thus, rats in the 10 s condition did not discriminate at either time point and showed no latency differences between the recent and remote retention tests. In contrast, for rats in the 60 s condition, a two-way repeated measures ANOVA revealed a significant main effect of recent versus remote test (F_(1, 17)_ = 5.55, *p =* .031), a significant main effect of context (F_(1, 17)_ = 7.10, *p =* .016), and no significant interaction (F_(1, 17)_ = 1.46, *p =* .24). Post hoc tests revealed a significant difference in aversive context latencies between recent and remote retention tests (*p =* .007) and a significant difference between aversive and neutral context latencies at the recent retention test (*p =* .023). Thus, rats in the 60 s group discriminated at the recent test and showed a decrease in retention from the recent to the remote test in the aversive context.

**Figure 2.**
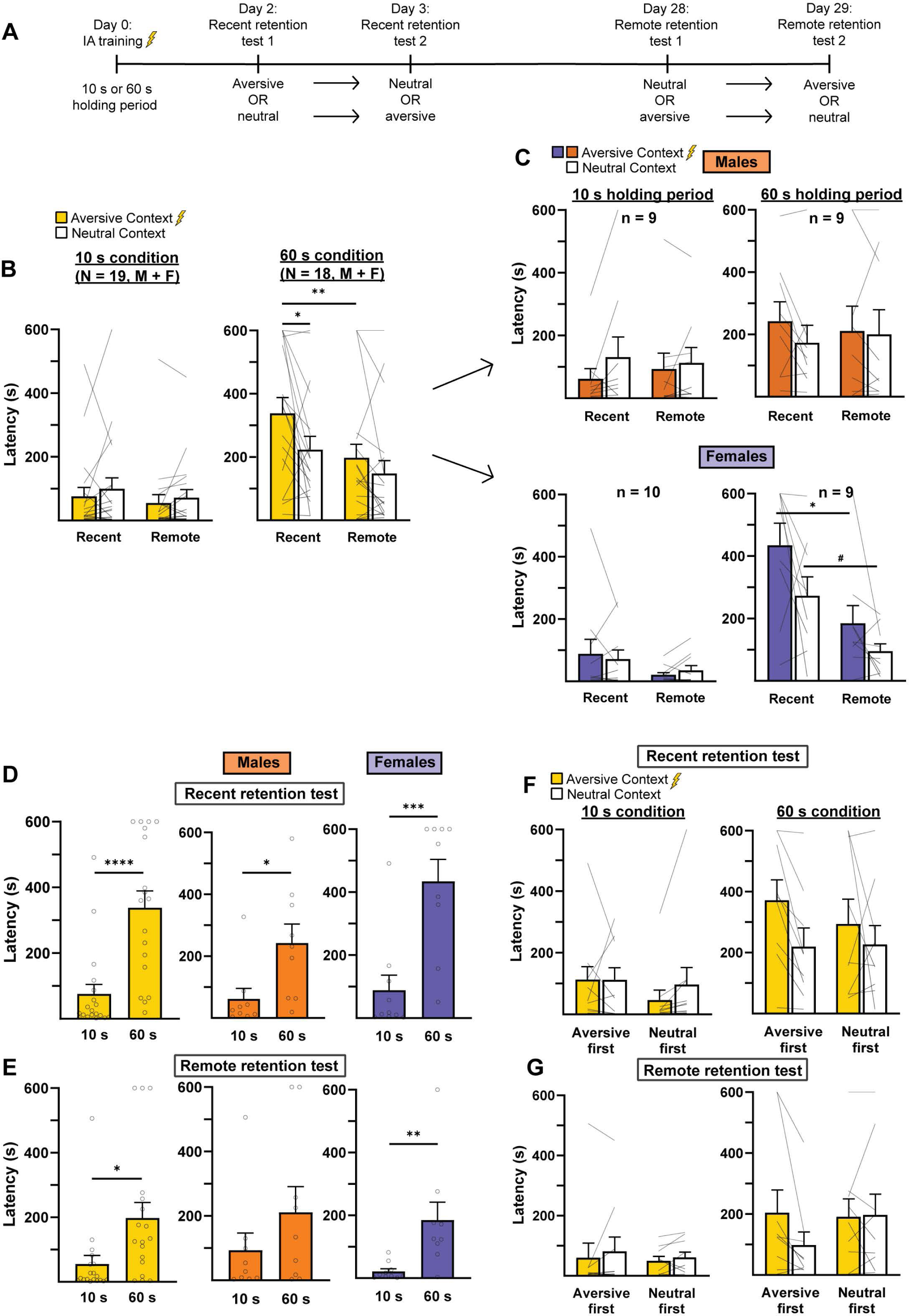
The 60 s holding period, compared to a 10 s holding period, increased memory strength at both recent and remote retention tests and promoted memory discrimination at the recent retention test. **A.** Experimental timeline. Rats underwent IA training while being retained for either 10 s or 60 s in the lit compartment before the door was retracted. Rats were then tested at recent and remote retention tests in each chamber. **B.** *Left,* Rats in the 10 s condition did not discriminate at either time point. *Right,* Rats in the 60 s condition discriminated at the recent retention test and generalized at the remote retention test, with a significant reduction in aversive context latencies (i.e., memory strength) between recent and remote retention tests. **C.** *Top left and right,* Male rats in the 10 s and 60 s conditions, respectively, did not discriminate at either time point. *Bottom left and right,* Female rats in either condition did not discriminate at either time point. However, female rats in the 60 s condition displayed a significant reduction in memory strength from the recent to the remote test, as well as a trend toward a reduction in neutral context latencies from the recent to remote test. **D.** *Left,* Rats had higher aversive context latencies (i.e., memory strength) in the 60 s condition compared to the 10 s condition at the recent retention test. Male (*middle*) and female (*right*) rats had higher aversive context latencies in the 60 s condition compared to the 10 s condition at the recent retention test. **E.** *Left,* Rats had higher aversive context latencies (i.e., memory strength) in the 60 s condition compared to the 10 s condition at the remote retention test. However, only female rats showed higher aversive context latencies in the 60 s condition compared to the 10 s condition at the remote test (*right*). **F.** Testing order had no effect on retention latencies in either holding condition at the recent retention test. **G.** Testing order had no effect on retention latencies in either holding condition at the remote retention test. * *p* < 0.05; ** *p* < 0.01; *** *p* < 0.001; **** *p* < 0.0001.

Figure 2C shows the data from Figure 2B split by sex (top: males; bottom: females). For male rats in the 10 s condition, a two-way repeated-measures ANOVA revealed no significant main effect of recent versus remote test (F_(1, 8)_ = 0.12, *p =* .74), a non-significant trend toward a main effect of context (F_(1, 8)_ = 5.11, *p =* .054), and no significant interaction (F_(1, 8)_ = 0.84, *p =* .39). Post hoc tests revealed no significant differences between conditions. Moreover, it appears that the rats had higher latencies in the neutral context compared to the aversive context. As this effect is in the opposite direction of what we would expect, this likely does not reflect a true difference. For female rats in the 10 s condition, a two-way repeated-measures ANOVA revealed a non-significant trend toward a main effect of recent versus remote test (F_(1, 9)_ = 3.76, *p =* .084), no significant main effect of context (F_(1, 9)_ = 0.01, *p =* .93), and no significant interaction (F_(1, 9)_ = 0.55, *p =* .48). Post hoc tests revealed no significant differences between conditions. Thus, when the data are split by sex, neither males nor females in the 10 s condition discriminated at either time point.

For the 60 s condition (Figure 2C, right), a two-way repeated-measures ANOVA of latencies for male rats revealed no significant main effect of recent versus remote test (F_(1, 8)_ = 0.001, *p =* .97), no significant main effect of context (F_(1, 8)_ = 3.05, *p =* .12), and no significant interaction (F_(1, 8)_ = 0.71, *p =* .42). In contrast, for female rats in the 60 s condition, a two-way repeated measures ANOVA revealed a significant main effect of recent versus remote test (F_(1, 8)_ = 16.29, *p =* .004), a non-significant trend toward a main effect of context (F_(1, 8)_ = 5.05, *p =* .055), and no significant interaction (F_(1, 8)_ = 0.69, *p =* .43). Post hoc tests revealed a significant difference in aversive context latency between recent and remote retention tests (*p =* .044) and a non-significant trend toward a difference in neutral context latencies between recent and remote retention tests (*p =* .053). Thus, when split by sex, neither males nor females discriminated at either time point. However, both groups still showed the same pattern of discrimination as when both sexes were combined. Additionally, only females showed a significant reduction in memory strength between recent and remote retention tests, suggesting that females drove the overall effect of testing time point.

Figure 2D shows how the 10 s versus 60 s conditions affected memory strength (i.e., aversive context latency) at the recent retention test. An unpaired t-test on the 10 s versus 60 s condition at the recent time point revealed that the 10 s rats had significantly poorer memory strength compared to the 60 s rats (t_(35)_ = 4.53, *p <* .0001). When split by sex, males and females both had significantly poorer memory strength in the 10 s condition compared to the 60 s condition at the recent time point (t_(17)_ = 2.56, *p =* .020 for males and t_(18)_ = 4.48, *p <* .001 for females). Thus, rats showed poorer memory strength in the 10 s condition compared to the 60 s condition at the recent retention test.

Figure 2E shows how the 10 s versus 60 s conditions affected memory strength at the remote retention test. An unpaired t-test on the two conditions at the remote time point again revealed that the 10 s rats had significantly poorer memory strength compared to the 60 s rats (t_(35)_ = 2.65, *p =* .012). When split by sex, males did not show any difference in memory strength between the 10 s and 60 s conditions at the remote retention test (t_(16)_ = 1.23, *p =* .24), whereas females again had significantly poorer memory strength in the 10 s condition compared to the 60 s condition (t_(17)_ = 2.96, *p =* .009).Thus, rats showed poorer memory strength in the 10 s condition compared to the 60 s condition at the remote time point, with females driving this effect.

Because the chamber testing order was counterbalanced, we wanted to determine whether testing order had an effect. Figure 2F shows the same retention test data as Figure 2B split by order of testing for the recent retention test. For rats in the 10 s condition at the recent time point, a two-way repeated measures ANOVA revealed no significant main effect of order (F_(1, 17)_ = 0.33, *p =* .57), no significant main effect of context (F_(1, 17)_ = 0.71, *p =* .41), and no significant interaction (F_(1, 17)_ = 1.01, *p* = .33). For rats in the 60 s condition at the recent time point, a two-way repeated measures ANOVA revealed no significant main effect of order (F_(1, 16)_ = 0.18, *p =* .68), a significant main effect of context (F_(1, 16)_ = 5.13, *p =* .038), and no significant interaction (F_(1, 16)_ = 0.78, *p* = .39). Post hoc tests revealed no significant differences between conditions. Thus, rats in both the 10 s and 60 s conditions did not show significant order effects at the recent retention test.

Figure 2G shows the remote retention test data split by order of testing. For the rats in the 10 s condition at the remote time point, a two-way repeated measures ANOVA revealed no significant main effect of order (F_(1, 17)_ = 0.09, *p =* .77), no significant main effect of context (F_(1, 17)_ = 1.61, *p =* .22) and no significant interaction (F_(1, 17)_ = 0.13, *p =* .72). For rats in the 60 s condition at the remote time point, a two-way repeated measures ANOVA revealed no significant main effect of order (F_(1, 16)_ = 0.25, *p =* .62), no significant main effect of context (F_(1, 16)_ = 2.62, *p =* .13), and a non-significant trend toward an interaction (F_(1, 16)_ = 3.28, *p =* .089). Post hoc tests revealed no significant differences between conditions. Thus, rats in both the 10 s and 60 s conditions did not show significant order effects at the remote retention test.

In order to investigate the possibility of reduced skewness in the 60 s condition compared to the 10 s condition, Pearson’s second coefficient of skewness was calculated for the recent retention test data for both conditions. For the 10 s data, the coefficient of skewness was 1.37, indicating a high degree of positive skewness, which is similar to what has been previously observed with IA retention latencies (Wahlstrom et al., 2018; Glickman et al., 2024). For the 60 s data, the coefficient of skewness was -0.34, indicating a very mild negative skew, likely reflecting the imposition of the artificial ceiling at 600 s. Thus, the 60 s holding condition resulted in reduced skewness compared to the 10 s condition. Based on these results, Experiments 2, 3 and 4 exclusively used the 60 s holding condition during training.

### Experiment 2

Experiment 2 used identical training procedures as Experiment 1 with the use of the 60 s holding condition, however rats were tested in both chambers on the same day for each retention test instead of being tested on two separate days. Figure 3A shows the experimental timeline. Figure 3B shows the recent and remote retention test latencies. A two-way repeated measures ANOVA revealed no significant main effect of recent versus remote test or interaction (F_(1, 18)_ = 0.33, *p =* .58; F_(1, 18)_ = 0.08, *p =* .78), but a significant main effect of context (F_(1, 18)_ = 9.47, *p =* .007). Post hoc tests revealed a non-significant trend toward a difference in aversive and neutral context latencies at the recent time point (*p =* .075). Thus, rats showed an overall difference in latencies in the aversive and neutral contexts, though they did not discriminate to a significant degree at either time point.

**Figure 3.**
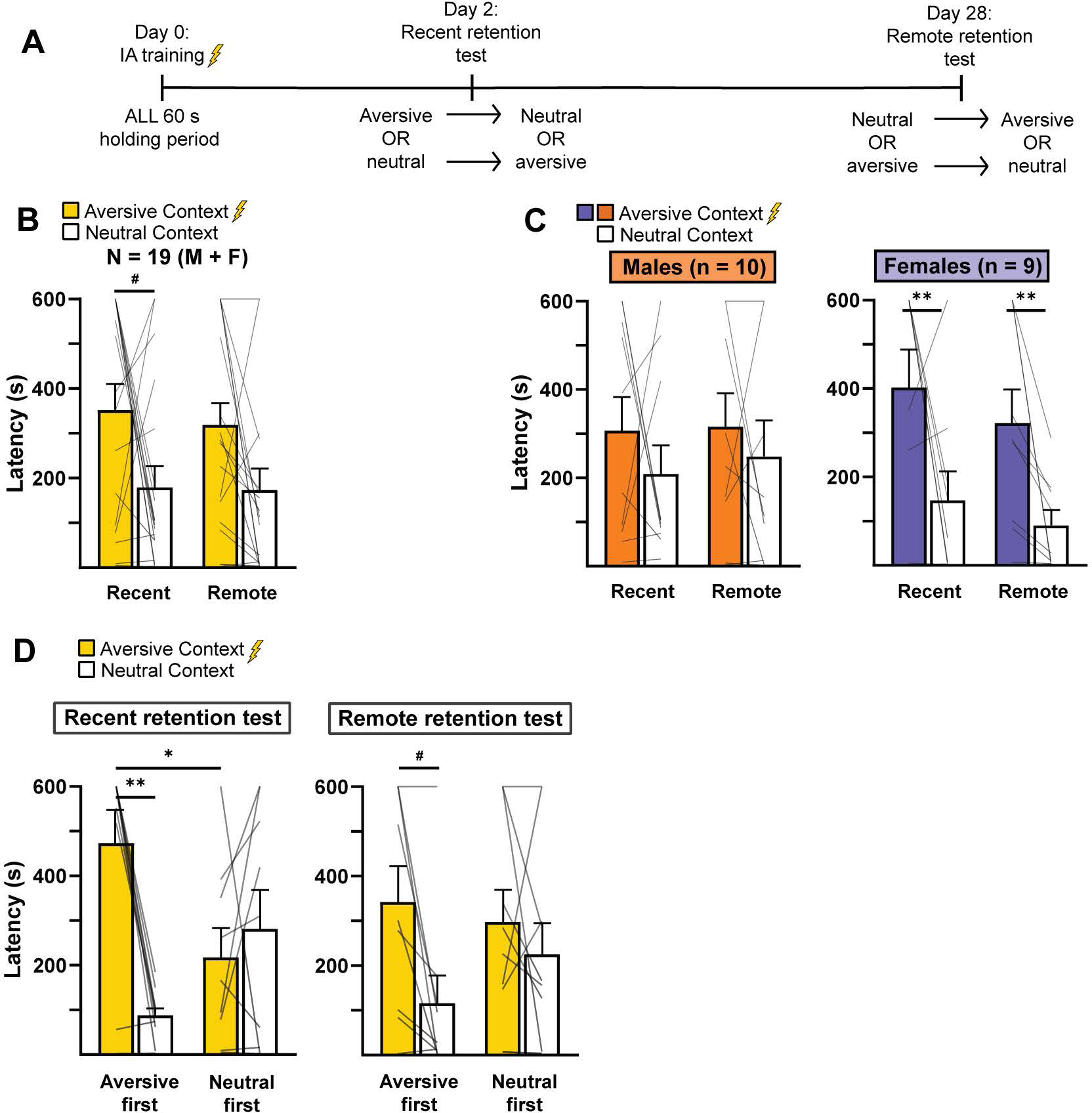
IA testing in both chambers on the same day resulted in significant testing order effects. **A.** Experimental timeline. Rats underwent IA training (all with the 60 s condition) and were then tested in each chamber on the same day at recent and remote time points. **B.** Rats demonstrated a non-significant trend toward discrimination in the recent retention test and did not discriminate at the remote retention test. **C.** *Left,* Male rats did not discriminate at either time point. *Right,* Female rats discriminated at both recent and remote time points. **D.** *Left,* At the recent retention test, rats discriminated when tested in the aversive chamber first and generalized when tested in the neutral chamber first. Additionally, rats tested in the aversive chamber first showed significantly higher aversive context latencies (i.e., memory strength) than those tested in the neutral chamber first. *Right,* At the remote retention test, rats tested in the aversive chamber first showed a trend toward discrimination that was not present in rats tested in the neutral chamber first. # *p* < 0.1; * *p* < 0.05; ** *p* < 0.01.

Figure 3C shows the same recent and remote retention test data split by sex (males on left, females on right). A two-way repeated-measures ANOVA for male rats revealed no significant main effect of recent versus remote test, context, or interaction (F_(1, 9)_ = 0.24, *p =* .64; F_(1, 9)_ = 2.41, *p =* .16; F_(1, 9)_ = 0.03, *p =* .86, respectively). A two-way repeated-measures ANOVA for female rats revealed no significant main effect of recent versus remote test or interaction (F_(1, 8)_ = 2.45, *p =* .16; F_(1, 8)_ = 0.11, *p =* .75, respectively), but a significant main effect of context (F_(1, 8)_ = 8.00, *p =* .022). Post hoc tests revealed that females showed a significant difference in aversive and neutral context latencies at both the recent time point (*p =* .004) and the remote time point (*p =* .005). Thus, when split by sex, male rats did not discriminate at either time point, whereas female rats discriminated at both time points.

Figure 3D shows the effects of order on these retention test results (recent retention test on left, remote retention test on right). A two-way repeated-measures ANOVA revealed that, at the recent retention test, there was no significant main effect of order (F_(1, 17)_ = 0.19, *p =* .67), a significant main effect of context (F_(1, 17)_ = 6.92, *p =* .018) and a significant interaction (F_(1, 17)_ = 13.53, *p =* .002). Post hoc tests revealed a significant difference in aversive context latencies between rats tested in the aversive context first versus neutral context first (*p =* .029) and a significant difference between aversive and neutral context latencies for rats tested in the aversive chamber first (*p* = .001). Thus, at the recent retention test, rats tested in the aversive chamber first discriminated, whereas rats tested in the neutral chamber first did not discriminate. Additionally, rats tested in the aversive chamber first showed significantly better memory strength than rats tested in the neutral chamber first. For the remote retention test data, a two-way repeated-measures ANOVA revealed no significant main effect of order or interaction (F_(1, 17)_ = 0.14, *p =* .71, F_(1, 17)_ = 1.76, *p =* .20), but a significant main effect of context F_(1, 17)_ = 6.56, *p =* .020). Post hoc tests revealed a non-significant trend toward a difference between aversive and neutral context latencies for rats tested in the aversive chamber first (*p =* .062). Thus, rats tested in the aversive chamber first showed a trend toward discrimination at the remote time point that was not present in the rats tested in the neutral chamber first. Therefore, due to the chamber testing order effects, subsequent experiments used the same testing procedures as Experiment 1.

### Experiment 3

Experiment 3 tested the effect of a pre-exposure to the neutral context. Figure 4A shows the experimental timeline. Figure 4B shows the recent and remote retention test latencies. A two-way repeated-measures ANOVA revealed no significant main effect of recent versus remote test (F_(1, 17)_ = 2.20, *p =* .16), no significant main effect of context (F_(1, 17)_ = 0.24, *p =* .63) and no significant interaction (F_(1, 17)_ = 0.22, *p =* .64). Thus, with a pre-exposure to the neutral chamber, rats generalized at both time points.

**Figure 4.**
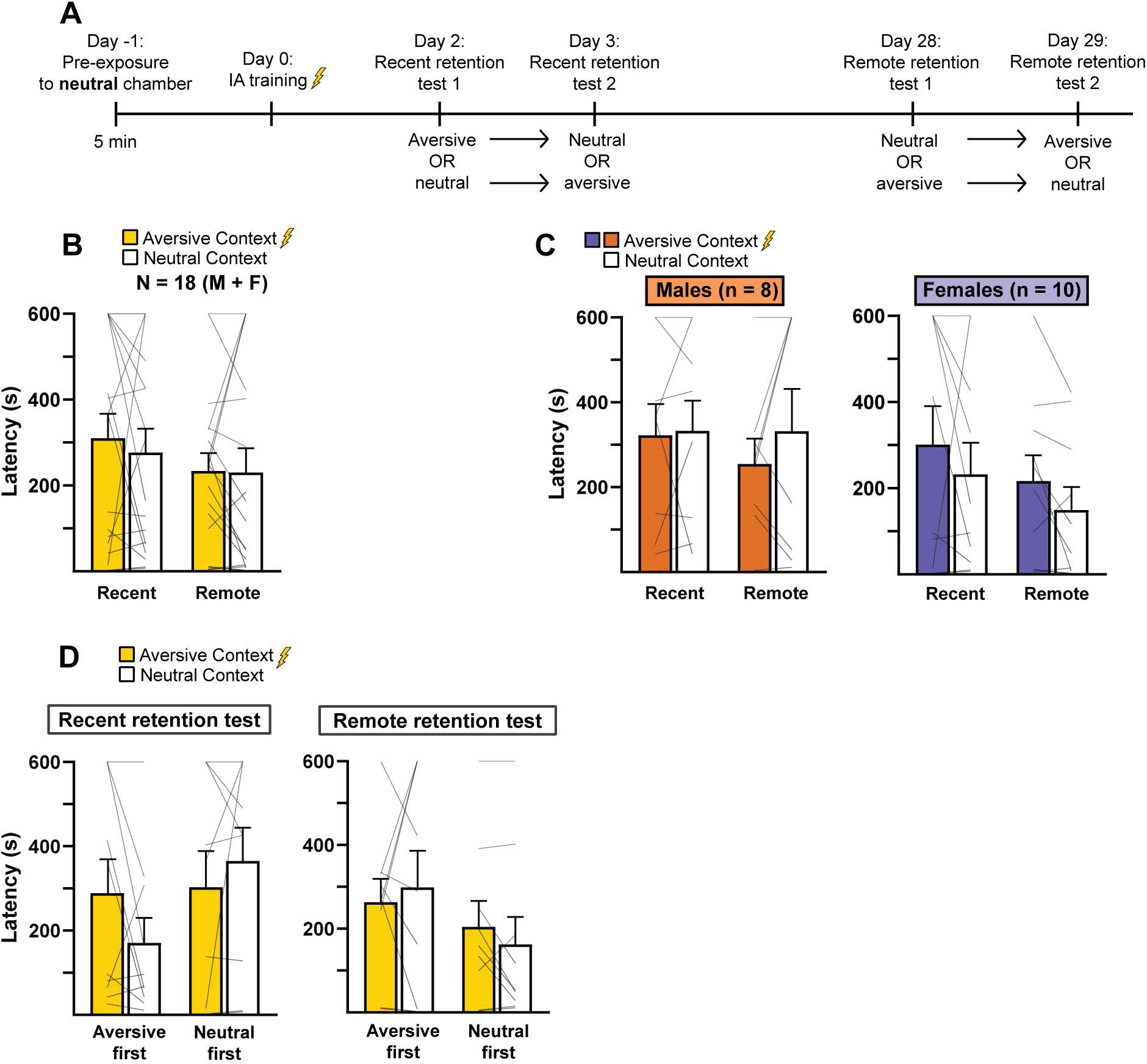
Pre-exposure to the neutral chamber promoted generalization at the recent retention test. **A.** Experimental timeline. Rats were given a 5 min pre-exposure to the neutral chamber, and the next day underwent IA training (all with the 60 s condition). Rats were then tested at recent and remote retention tests in each chamber. **B.** Rats generalized at both recent and remote retention tests. **C.** Males (*left*) and females (*right*) generalized at both time points. **D.** Testing order had no effect on retention latencies at both recent (*left*) and remote (*right*) retention tests.

Figure 4C shows the same recent and remote retention test data split by sex (males on left, females on right). A two-way repeated-measures ANOVA for the males revealed no significant main effect of recent versus remote test (F_(1, 7)_ = 0.37, *p =* .56), no significant main effect of context (F_(1, 7)_ = 0.71, *p =* .43) and no significant interaction (F_(1, 7)_ = 0.54, *p =* .49). Similarly, for females, a two-way repeated-measures ANOVA revealed no significant main effect of recent versus remote test (F_(1, 9)_ = 1.83, *p =* .21), no significant main effect of context (F_(1, 9)_ = 1.93, *p =* .20), and no significant interaction (F_(1, 9)_ < 0.01, *p =* .99). Thus, when split by sex, both males and females generalized at both time points.

Figure 4D shows the effects of order on these retention test results (recent retention test on left, remote retention test on right). A two-way repeated-measures ANOVA revealed that, at the recent retention test, there was no main effect of order (F_(1, 17)_ = 1.13, *p =* .30), no main effect of context (F_(1, 17)_ = 0.30, *p =* .59), and a non-significant trend toward an interaction (F_(1, 17)_ = 3.20, *p =* .092). Post hoc tests, however, revealed no significant differences between conditions. For the remote retention test data, a two-way repeated-measures ANOVA revealed no significant main effect of order (F_(1, 16)_ = 1.13, *p =* .30), no significant main effect of context (F_(1, 16)_ < 0.01, *p =* .94) and no significant interaction (F_(1, 16)_ = 0.90, *p =* .36). Thus, testing order did not significantly alter the results, with the exception of the trend toward an interaction at the recent time point.

### Experiment 4

Experiment 4 tested the effect of a pre-exposure to the aversive context. Figure 5A shows the experimental timeline. Figure 5B shows the recent and remote retention test latencies. A two-way repeated measures ANOVA revealed no significant main effect of recent versus remote test (F_(1, 17)_ = 2.46, *p =* .14), a significant main effect of context (F_(1, 17)_ = 54.03, *p <* .001), and a significant interaction (F_(1, 17)_ = 37.92, *p <* .001). Post hoc tests revealed a significant difference in aversive context latencies between the recent and remote retention tests (*p <* .001), a significant difference between aversive and neutral context latencies at the recent retention test (*p <* .001) and a significant difference between aversive and neutral context latencies at the remote retention test (*p <* .001). Thus, with a pre-exposure to the aversive chamber, rats discriminated at both time points, albeit with a significant reduction in memory strength between the recent and remote time point.

**Figure 5.**
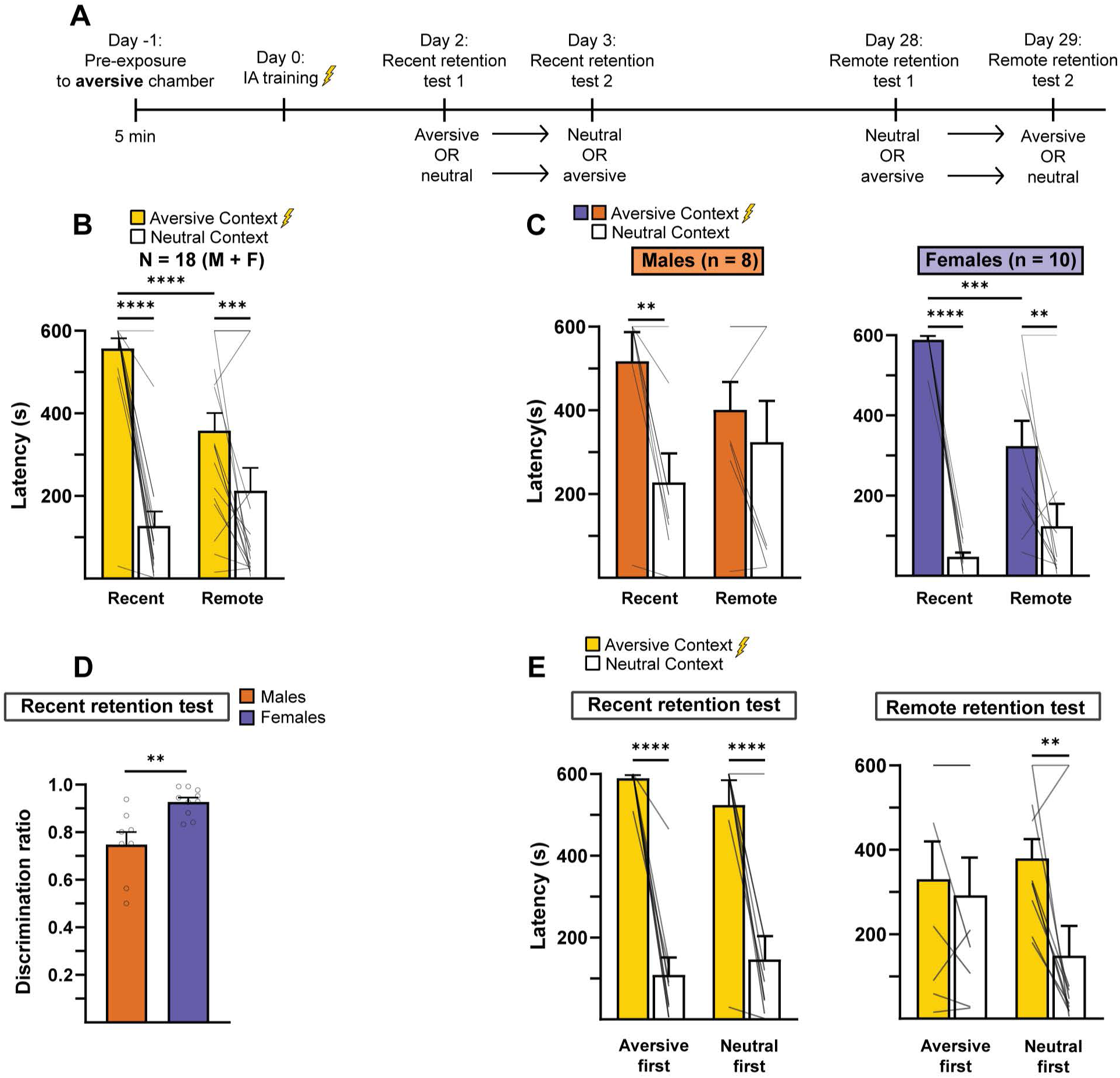
Pre-exposure to the aversive chamber promoted discrimination and memory strength at both recent and remote retention tests. **A.** Experimental timeline. Rats were given a 5 min pre-exposure to the aversive chamber, and the next day underwent IA training (all with the 60 s condition). Rats were then tested at recent and remote retention tests in each chamber. **B.** Rats discriminated at both recent and remote retention tests, with a significant reduction in aversive context latencies (i.e., memory strength) between recent and remote time points. **C.** *Left,* Males discriminated at the recent retention test and generalized at the remote retention test. *Right,* Females discriminated at both recent remote retention tests, with a significant reduction in aversive context latencies (i.e., memory strength) between recent and remote time points. **D.** *Top,* Discrimination ratio calculation. *Bottom,* Discrimination ratios calculated for males and females indicated enhanced discrimination in females at the recent retention test compared to males. **E.** *Left,* Testing order had no effect on retention latencies at the recent retention test. *Right,* At the remote retention test, rats tested in the aversive chamber first generalized, whereas rats tested in the neutral chamber first discriminated. * *p* < 0.05; ** *p* < 0.01; *** *p* < 0.001; **** p < 0.0001.

Figure 5C shows the same retention test data split by sex (males on left, females on right). A two-way repeated-measures ANOVA for the males revealed no significant main effect of recent versus remote test (F_(1, 7)_ = 0.04, *p =* .85), a significant main effect of context (F_(1, 7)_ = 11.28, *p =* .012), and a significant interaction (F_(1, 7)_ = 10.75, *p =* .014). Post hoc tests revealed a significant difference between aversive and neutral context latencies at the recent retention test (*p =* .002), but no difference between aversive and neutral context latencies at the remote retention test.

Thus, males discriminated at the recent time point, but not at the remote time point. For females, a two-way repeated-measures ANOVA revealed a non-significant trend toward a main effect of recent versus remote test (F_(1, 9)_ = 3.57, *p =* .091), a significant main effect of context (F_(1, 9)_ = 85.21, *p <* .001), and a significant interaction (F_(1, 9)_ = 30.60, *p <* .001). Post hoc tests revealed a significant difference in aversive context latencies between the recent and remote retention tests (*p <* .001), a significant difference between aversive and neutral context latencies at the recent retention test (*p <* .001) and a significant difference between aversive and neutral context latencies at the remote retention test (*p =* .003). Thus, females discriminated at both time points and showed a significant reduction in memory strength between the recent and remote time point.

Even though both males and females discriminated at the recent retention test, the data suggested a difference in the degree of discrimination. To investigate this possibility, discrimination ratios [aversive context latency/(aversive context latency + neutral context latency)] were calculated.

As shown in Figure 5D, an unpaired t-test revealed that females showed significantly greater discrimination ratios than males (t_(16)_ = 3.52, *p =* .003). Thus, at the recent time point, females discriminated between aversive and neutral contexts to a greater extent than males.

Figure 5E shows the effects of testing order on these retention test results (recent retention test on left, remote retention test on right). A two-way repeated-measures ANOVA revealed that, at the recent retention test, there was no significant main effect of order (F_(1, 16)_ = 0.06, *p =* .81), a significant main effect of context (F_(1, 16)_ = 95.72, *p <* .001), and no significant interaction (F_(1, 16)_ = 1.40, *p =* .26). Post hoc tests revealed a significant difference between aversive and neutral context latencies for rats tested in both the aversive first (*p <* .001) and neutral first (*p <* .001) conditions. For the remote retention test data, a two-way repeated measures ANOVA revealed no main effect of order (F_(1, 16)_ = 0.22, *p =* .65) a main effect of context (F_(1, 16)_ = 10.42, *p =* .005) and a significant interaction (F_(1, 16)_ = 5.31, *p =* .035). Post hoc tests revealed a significant difference between aversive and neutral context latencies for rats tested in the neutral chamber first (p = .003), but not for rats tested in the aversive chamber first. Thus, there were no significant order effects at the recent retention test, but there were significant order effects at the remote retention test, as only rats tested in the neutral chamber first discriminated.

### Ancillary analyses

In addition to the analyses in Experiments 1-4, two additional analyses were conducted. First, to determine whether there were differences in memory strength at the recent retention test between the two pre-exposure groups (Experiments 2 and 3) and the no pre-exposure group from Experiment 1 (60 s condition only), the aversive context latencies were compared across all three groups. Figure 6A shows the results from comparing aversive context latencies from the recent retention tests in the no pre-exposure group, neutral context pre-exposure group, and aversive context pre-exposure group. A one-way between-subjects ANOVA revealed a significant effect of testing condition on retention latency (F_(2, 52)_ = 8.32, *p* < .001). Post hoc tests revealed a significant difference between the no pre-exposure group and the aversive context pre-exposure group (*p* = .007), a significant difference between the neutral context pre-exposure group and the aversive context pre-exposure group (*p* = .001), and no significant difference between the no pre-exposure group and neutral context pre-exposure group. Thus, the aversive context pre-exposure significantly increased memory strength compared to the no pre-exposure condition and the neutral pre-exposure condition.

**Figure 6.**
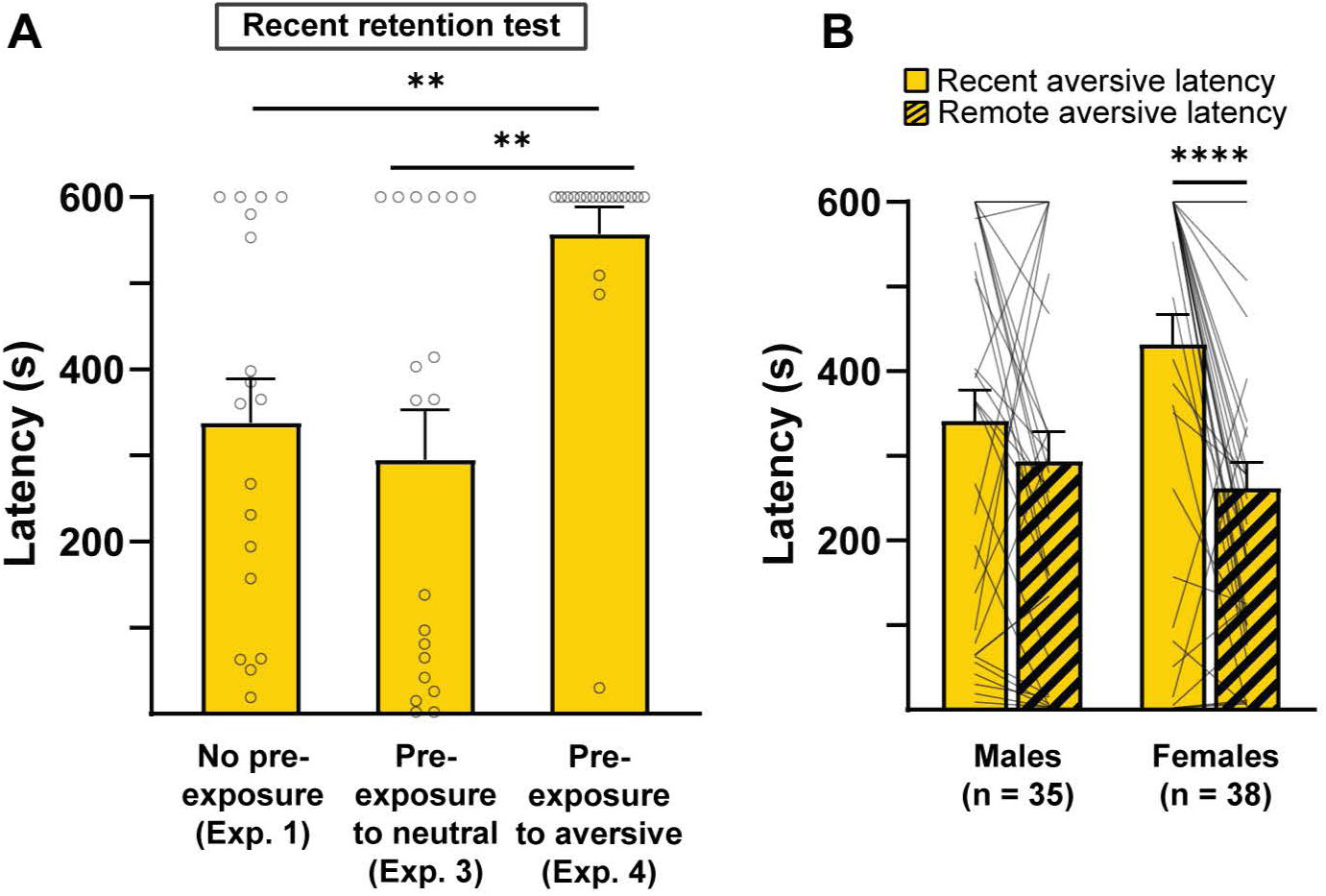
**A.** Recent aversive retention latency (i.e., memory strength) was increased in the aversive context pre-exposure condition compared to the no pre-exposure condition (60 s condition only) and the neutral context pre-exposure condition, whereas memory strength in the neutral context pre-exposure condition did not differ from that of the no pre-exposure condition. **B.** When data from all experiments were combined, females showed a significant reduction in memory strength between the recent and remote retention tests that was not seen in males. ** *p* < 0.01; **** p < 0.0001.

Second, as there appeared to be a pattern across the experiments in which females, but not males, showed a reduction in memory strength from the recent to the remote tests, the data from all experiments (Experiments 1-4, 60 s condition only from Experiment 1) were combined and analyzed between the early and late tests. Figure 6B shows the results from combining recent and remote aversive context retention latencies across all conditions (Experiments 1-4) for males and females. A two-way repeated measures ANOVA revealed a significant main effect of recent versus remote test (F_(1, 71)_ = 18.04, *p* < .001), no significant main effect of sex (F_(1, 71)_ = 0.41, *p* = .53), and a significant interaction (F_(1, 71)_ = 5.75, *p* = .019). Post hoc tests revealed a significant difference between recent and remote retention latencies for females (*p* < .001) but not for males. Thus, these results confirmed the overall pattern of females, but not males, showing a reduction in memory strength from the recent to the remote retention test.

## Discussion

The present findings point to several different procedures capable of biasing IA to produce changes in memory strength and/or precision, as assessed at recent and remote time points. The results indicate that keeping rats in the lit compartment of the aversive chamber for 60 s on training day, as opposed to 10 s, enhanced discrimination at the recent retention test and improved memory strength at both recent and remote retention tests. The present findings also suggest that testing rats in both chambers on the same day produced testing order effects. The last set of experiments then examined how pre-exposures to the neutral and aversive chambers, respectively, influenced memory strength and precision. The results indicate that pre-exposure to the neutral chamber promoted generalization without affecting memory strength, whereas pre-exposure to the aversive chamber promoted discrimination along with improved memory strength. Finally, the current work observed sex differences in several of the experiments, with the overall pattern suggesting enhanced discrimination in females compared to males.

### 10 s versus 60 s holding conditions

Prior work, including from our laboratory, has typically used IA procedures wherein rats are kept in the lit compartment for only 10 s during training or even are placed in the lit compartment with the door already retracted (Lalumiere et al., 2004; Lalumiere and McGaugh, 2005; Glickman et al., 2024). Such procedures frequently lead to positive skewing, in which many rats display low retention latencies, even as some display much higher retention latencies (Wahlstrom et al., 2018; Glickman et al., 2024). Therefore, we hypothesized that additional time to encode the context before the footshock would improve memory strength and reduce the degree of skew, an idea that the findings of Experiment 1 support. Thus, the remaining experiments exclusively used the 60 s holding condition.

### Context pre-exposure

Prior work suggests that pre-exposure to different contexts alter the resulting memory strength and/or discrimination (Keiser et al., 2017). Therefore, the present experiments attempted to identify procedures that might bias rats toward generalization versus discrimination in IA. One possible explanation for the relatively weak discrimination at the recent retention test in Experiment 1 was that the rats did not have prior exposure to the neutral chamber and, therefore, perceived the neutral chamber as closely related to the aversive chamber. Thus, we hypothesized that giving rats a separate exposure to the neutral chamber without a footshock would facilitate encoding of the neutral chamber, thereby enhancing contextual discrimination during the retention test. However, the results suggest just the opposite, as pre-exposure to the neutral chamber promoted generalization at both the recent and remote test. Previous work found that pre-exposure to the neutral context 1 min before IA training promoted generalization, whereas pre-exposure 2 min before training restored strong discrimination (Atucha and Roozendaal, 2015). However, it is unclear how those findings fit with the current results. Atucha and Roozendaal used Sprague-Dawley rats in contrast to the present work using Long-Evans rats, and prior studies have observed strain differences in rats in various learning and memory tasks (Tonkiss et al., 1992; Andrews et al., 1995; Turner and Burne, 2014; Olivera-Pasilio and Dabrowska, 2023). Moreover, the present study used a pre-exposure 1 d in advance, in contrast to the Atucha and Roozendaal study, providing time for consolidation of the contextual memory over a full day prior to training.

Notably, unpublished work from one author (JJR) found that pre-exposure to the neutral chamber promotes generalization even when the pre-exposure occurs 7 d before training. Together with the present results, these findings suggest that generalization does not simply result from temporal proximity between pre-exposure and training. One possibility is that the context pre-exposure establishes a memory trace for the neutral chamber that, due to contextual similarity, is reactivated during training. Consequently, when the footshock occurs, the contextual trace associated with the footshock encompasses both contexts. Future work using approaches such as neuronal ensemble tagging techniques (for review, see Josselyn and Tonegawa, 2020) may be helpful for examining this issue. Experiment 3 also found that the pre-exposure to the neutral chamber appeared to promote generalization without promoting memory strength when compared to the results of Experiment 1. This finding supports the idea that memory precision and memory strength are dissociable attributes of memory. However, the neurobiological mechanisms that separately influence these processes have not been well elucidated.

In contrast to neutral context pre-exposure, Experiment 4 found that pre-exposure to the aversive context enhanced memory strength. This finding is consistent with a large body of evidence indicating that such pre-exposure improves memory strength in contextual fear conditioning (Fanselow, 1990; Bae et al., 2015). Moreover, aversive context pre-exposure also promoted discrimination, even at the remote time point. However, the effect at the remote time point appears to have been primarily driven by the females. In this case, perhaps the reason for the improvement in memory strength is that, similar to Experiment 3, rats established a memory trace for the context that they were pre-exposed to. However, because the pre-exposure context was the same as the aversive context, this memory trace was reinforced with training, which promoted memory strength. The enhanced discrimination seen in Experiment 4 is likely related to the overall increase in memory strength, with perhaps a small decrease in neutral context latencies also contributing to discrimination.

### Sex differences in memory strength and precision

The present results identified a pattern of females showing enhanced discrimination compared to males, particularly in Experiment 4. An additional sex difference was observed in the tendency for females to show a reduction in memory strength between the recent end remote time point, whereas males displayed a relatively consistent amount of memory strength across time.

Previous work has demonstrated a tendency for episodic memories to lose their strength from recent to remote time points (Asok et al., 2018), but the current study found evidence of this effect only in females. Past work has found evidence for sex differences in contextual fear conditioning, with males typically showing better retention than females (Maren et al., 1994; Russo and Parsons, 2021) and females showing elevated levels of generalization compared to males (Keiser et al., 2017). Intriguingly, sex differences in contextual fear conditioning can be altered depending on several behavioral factors, such as context pre-exposure (Asok et al., 2019; Trott et al., 2022) and the interval between placement in the chamber and shock during training (Wiltgen et al., 2001). However, in contrast to prior work (Keiser et al., 2017), the present study found that females tended to discriminate more than males, with a pattern of increased memory strength compared to males. One possible explanation for the conflicting results is that IA measures retention latencies, as opposed to freezing, which is typically used in contextual fear conditioning experiments. Prior evidence suggests that females may display non-freezing behavior that is still indicative of retention of fear memory. For example, darting behaviors in auditory fear conditioning studies have been observed in females particularly (Gruene et al., 2015; Mitchell et al., 2022), indicating that freezing alone may not be the most reflective measure of retention in rodents. Thus, latencies, which do not depend exclusively on freezing, may provide a larger “umbrella” measure of retention for both males and females.

## Conclusion

Together, the present results found that 1) keeping rats in the lit compartment on training day for 60 s, relative to 10 s, resulted in improved discrimination at the recent retention test and improved memory strength at both the recent and remote retention tests, 2) testing rats in each IA chamber on the same day led to testing order effects, 3) a pre-exposure to the neutral chamber promoted generalization, and 4) a pre-exposure to the aversive chamber promoted discrimination and memory strength. These procedures provide a roadmap for investigating questions about memory strength and precision and point to critical experiments for understanding the effects of different types of context pre-exposure on contextual encoding. In addition, the current work provides evidence for how differences in behavioral procedures at the time of encoding can lead to different results in retention.

## Acknowledgements

This work was supported by the National Institutes of Health grant MH132223 (JJR and RTL). The authors would like to thank Matthew McGregor, Alexa Zimbelman, Hanxiao Liu, Kaitlyn Jones, and Chris Nguyen for their helpful comments on a draft of the manuscript.

